# Integrative Multi-Omics Analysis of IFNγ-induced Macrophages and Atherosclerotic Plaques Reveals Macrophage-dependent STAT1-Driven Transcription in Atherosclerosis

**DOI:** 10.1101/2024.09.06.611606

**Authors:** Mahdi Eskandarian Boroujeni, Natalia Lopacinska, Aleksandra Antonczyk, Katarzyna Kluzek, Joanna Wesoly, Hans AR Bluyssen

**Affiliations:** Human Molecular Genetics Research Unit, Institute of Molecular Biology and Biotechnology, Faculty of Biology, Adam Mickiewicz University, Poznań, Poland; Laboratory of High-Throughput Technologies, Institute of Molecular Biology and Biotechnology, Faculty of Biology, Adam Mickiewicz University, Poznań, Poland

**Keywords:** atherosclerosis, IFNγ signaling, STAT1, multi-omics integration, macrophages, single cell RNA-seq

## Abstract

This study investigates the role of STAT1-mediated IFNγ signaling in atherosclerosis progression through multi-omics integration and analysis of human and mouse models of atherosclerotic lesions. By integrating ATAC-seq, ChIP-seq, and RNA-seq data from IFNγ-treated bone marrow-derived macrophages, we identified 1139 STAT1-dependent integrative genes that show chromatin accessibility, differential epigenetic marks (H3K27ac, H3K4me1, H3K4me3), prominent transcription factor binding patterns (STAT1 and PU.1), and active transcription. These genes were also enriched for lipid metabolism and atherosclerosis-related pathways. We then validated our findings by tracing the expression of these genes in human atherosclerotic lesions and in ApoE-/- and LDLr-/-mouse models, revealing significant correlations with LDL cholesterol and diseased vessel traits. Single-cell RNA-seq of human and mouse atherosclerotic samples showed dynamic changes in macrophage subtypes, with foamy and tissue-resident macrophages displaying increased STAT1 activity. This comprehensive multi-omics approach provides new insights into the transcriptional regulation of atherosclerosis progression mediated by STAT1-PU.1 co-binding and IFNγ signaling. Moreover, our data delineates a STAT1-dependent gene signature, highlighting the potential of these integrative genes as biomarkers and therapeutic targets in atherosclerosis.

## Introduction

The development and progression of atherosclerosis, a chronic inflammatory disease of the arterial wall, involves complex interactions between various cell types and signaling pathways. Among these, the role of macrophages and interferon-gamma (IFNγ) signaling has emerged as a critical factor in atherosclerotic lesion formation and progression (Hansson & Hermansson, 2011; Libby et al., 2019; Ramji & Davies, 2015).

Recent advances in multi-omics technologies have provided unprecedented opportunities to investigate the molecular mechanisms underlying atherosclerosis at various levels of biological organization. By integrating data from chromatin accessibility assays, epigenetic modifications, transcription factor binding patterns, and gene expression profiles, researchers can now gain a more comprehensive understanding of the regulatory networks driving disease progression (Doran et al., 2021; Hasin, Seldin, & Lusis, 2017).

One key player in the IFNγ signaling pathway is the transcription factor Signal Transducer and Activator of Transcription 1 (STAT1). STAT1 is activated by IFNγ and plays a crucial role in mediating its downstream effects, including the regulation of genes involved in inflammation, lipid metabolism, and immune responses (Chmielewski et al., 2014; Platanitis & Decker, 2018). However, the precise mechanisms by which STAT1-mediated IFNγ signaling contributes to atherosclerosis progression remain incompletely understood.

Macrophages play a central role in atherosclerosis, from early foam cell formation to advanced plaque development and potential rupture. These cells exhibit remarkable plasticity, adopting various phenotypes in response to environmental cues within the atherosclerotic lesion. The heterogeneity of macrophage populations in atherosclerotic plaques has been increasingly recognized, with distinct subsets showing pro-inflammatory, anti-inflammatory, or lipid-handling properties (Gui, Zheng, & Cao, 2022; Hou et al., 2023).

The interplay between macrophages and IFNγ signaling is particularly relevant in the context of atherosclerosis. IFNγ, primarily produced by T cells and natural killer cells, can profoundly influence macrophage function, promoting a pro-inflammatory phenotype and enhancing lipid uptake. Moreover, STAT1 activation in macrophages has been shown to exacerbate atherosclerosis in animal models, suggesting a critical role for this signaling axis in disease progression (Agrawal et al., 2007; Sikorski, Wesoly, & Bluyssen, 2014; Voloshyna, Littlefield, & Reiss, 2014).

Epigenetic regulation, namely histone modifications and chromatin remodeling, has emerged as a chief mechanism controlling macrophage responses in atherosclerosis. Recent studies have demonstrated that alterations in histone acetylation and methylation patterns can significantly impact macrophage activation states and their contribution to plaque development (Bekkering et al., 2014; Kuznetsova et al., 2020). Understanding how IFNγ and STAT1 signaling intersect with these epigenetic processes could provide new insights into the molecular basis of atherosclerosis This study aims to elucidate the role of STAT1-mediated IFNγ signaling in atherosclerosis through a comprehensive multi-omics approach. By integrating ATACseq, ChIP-seq, and RNA-seq data from IFNγ-treated bone marrow-derived macrophages, we identified a set of STAT1-dependent integrative genes that exhibit specific epigenetic and transcriptional characteristics. These genes were further investigated in human atherosclerotic lesions and mouse models of atherosclerosis to validate their relevance to disease progression. Additionally, we used public single-cell RNA sequencing data to examine the dynamic changes in macrophage subtypes within atherosclerotic plaques, focusing on the expression patterns of STAT1-dependent genes in different macrophage populations. This approach allows for a more nuanced understanding of the cellular heterogeneity within atherosclerotic lesions and the specific contributions of different macrophage subtypes to disease progression. By combining these multi-omics approaches with in-depth bioinformatic analyses, our study aims to provide new insights into the transcriptional regulation of atherosclerosis mediated by STAT1 and IFNγ signaling. The identification of a STAT1-dependent gene signature may not only enhance our understanding of the disease mechanisms but also highlight potential biomarkers and therapeutic targets for atherosclerosis.

## Results

### Multi-Omics based integration of Macrophage IFNγ-stimulated transcriptional changes: Identification of IFNγ-responsive STAT1-dependent integrated genes

We sought to identify IFNγ-responsive STAT1-dependent integrated genes in macrophages with the implications in atherosclerosis (Figure 1). In this regard, we focused on bone marrow-derived macrophages that were treated with IFNγ at a defined time point (approximately 2h). To gain a deeper understanding of IFNγ-responsive STAT1-dependent transcriptional regulation in macrophages at different levels, we collected the datasets that examined chromatin accessibility (Platanitis et al., 2022) and acetylation (H3K27ac) (Piccolo et al., 2017) and methylation (H3K4me1 and H3K4me3) (Ostuni et al., 2013) along with the binding pattern of key transcription factors STAT1 (Piccolo et al., 2017; Platanitis et al., 2019) and PU.1 (Langlais, Barreiro, & Gros, 2016) upon IFNγ treatment of mouse macrophages (Table S1). To minimize batch effects, we prioritized selecting datasets that contained a greater number of markers and were, whenever feasible, conducted by a single laboratory, followed by normalizing all the samples as described in materials and methods. We also conducted our in-house transcription profiling of IFNγ-treated macrophages at specified time points (0, 0.5, 2, 4, 8, 24) to cover a wider range of time points (Table S2). To obtain a IFNγ-responsive list of STAT1-dependent integrative genes that meet all these criteria 1) are accessible at the chromatin level, 2) show differential epigenetic marks, 3) have a clear STAT1 binding site and a detectable co-binding with PU.1 as a lineage-determining transcription factor 4) actively being transcribed, these ATAC-seq, ChIP-seq and RNA-seq were integrated at these different levels. Subsequently, we decided to trace the expression pattern of these integrative genes in human patients with atherosclerotic lesions and in two most frequently used mouse models for atherosclerosis ApoE-/-mice and LDLr-/-mice (Table S1). This led to the identification of STAT1-dependent gene signature that distinctly contributes to the atherosclerosis progression.

**Figure 1.**
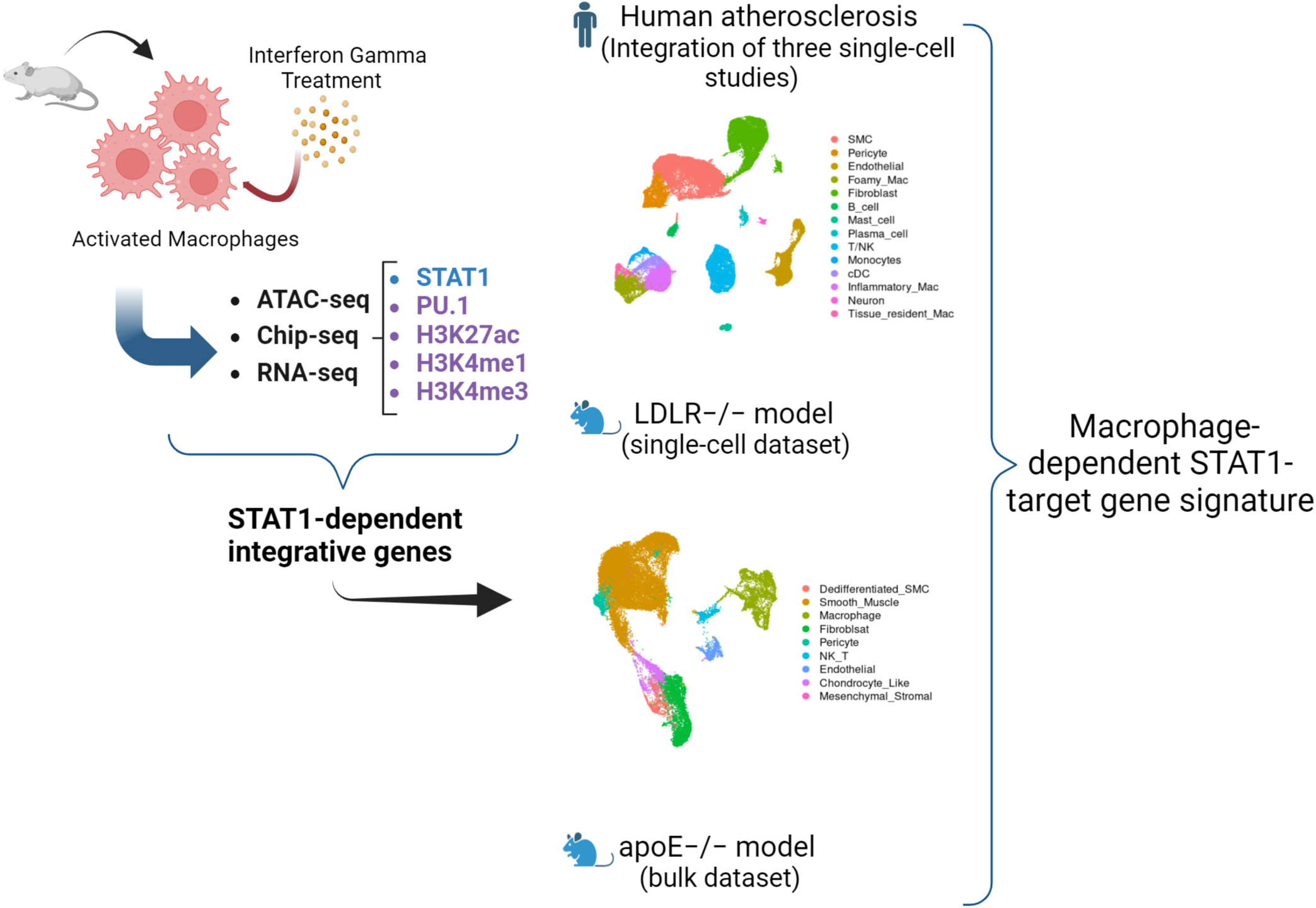
The schematic overview of the study. To identify IFNγ-responsive STAT1-dependent integrated genes in macrophages, activated macrophages are treated with interferon gamma, followed by integration of ATAC-seq, ChIP-seq, and RNA-seq datasets (See Table S1 for sample details). The identified STAT1-dependent integrative genes are then traced in human patients with atherosclerotic lesions and in two most frequently used mouse models for atherosclerosis ApoE-/- mice and LDLr-/- mice. This led to the identification of macrophage-dependent STAT1-target gene signatures across different models of atherosclerosis.

In our analysis of ATAC-seq and ChIP-seq datasets, after merging all the peaks, we were interested to assess how these peaks are correlated across various datasets (Figure 2A). To obtain a global view, we first examined the pairwise correlation of the peaks derived from all genomic regions and then we zoomed in only those peaks that are restricted to the promoter region (−3000, 3000). As shown in figure 2A, the correlation pattern looks similar regardless of the location of the peaks (all genomic regions vs promoter-exclusive regions). Moreover, we observed a strong correlation of STAT1 with H3K27ac and H3K4me1 in IFNγ-treated samples, indicating the association of STAT1 binding with these epigenetic marks upon IFNγ exposure.

**Figure 2.**
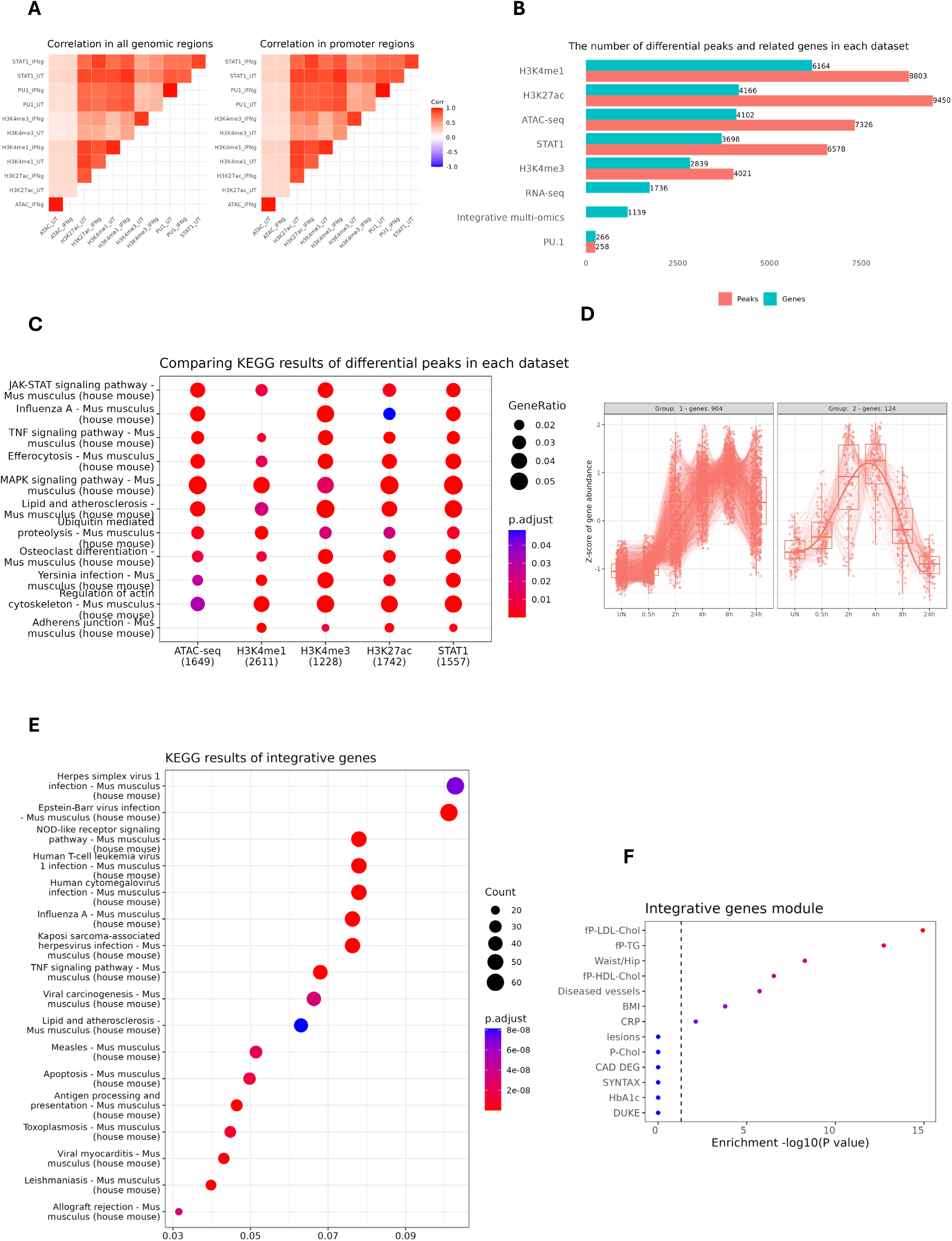
Identification and characterization of IFNγ-responsive STAT1-dependent integrated genes. (A) the pairwise correlation of the peaks derived from all genomic regions (left) and the peaks that are restricted to the promoter region (−3000, 3000). (B) The horizontal bar plot showing the number of differential peaks and their related genes (entrez gene id) in each dataset. For ATAC-seq, H3K4me1 & H3K4me3, a fold change cut-off of 2 was selected, whereas for PU.1, STAT1 and H3K27ac, a cut-off of 4 was set. The bars are sorted based on the number of genes in each dataset. (C) KEGG pathway analysis of differential peaks for each dataset. The numbers in the parentheses indicate the total number of annotated genes in each dataset. (D) The expression pattern of STAT1-dependent integrative genes from our RNA-seq experiment performed on IFNγ-treated mouse macrophages at different time points. (E) KEGG pathway analysis of STAT1-dependent integrative genes. (F) The clinically phenotypic enrichment of integrative gene module using STARNET database. The color intensity represents the statistical significance of observed traits.

To ensure our integrative genes are IFNγ-responsive, we performed differential peak analysis on normalized ATAC-seq and ChIP-seq datasets. Then, the differentially enriched regions were integrated with transcriptionally upregulated genes under IFNγ treatment. The multi-omics integration resulted in 1139 genes (Figure 2B; Table S3). Subsequently, we looked at the expression pattern of these 1139 genes in our time-series RNA-seq experiment, and we found two major gene expression patterns indicating progressively increasing expression levels at late time points from 2 to 24h (Figure 2D). The KEGG analysis of differential peaks in each dataset (except PU.1) highlighted relevant terms including JAK-STAT, MAPK and TNF signaling pathways and lipid and atherosclerosis (Figure 2C). It is noteworthy that PU.1 binding is not remarkably affected by IFNγ treatment, since we detected a low number of differential peaks in comparison with STAT1 and other epigenetic marks and there is no enriched KEGG terms with respect to differentially PU.1 binding. As for integrative genes, the KEGG analysis showed enrichment for lipid and atherosclerosis (Figure 2E). Moreover, we queried the Stockholm-Tartu Atherosclerosis Reverse Network Engineering Task (STARNET) gene regulatory networks across seven cardiometabolic tissues (Koplev et al., 2022). We observed that our integrative genes were highly associated with phenotypic traits such as LDL cholesterol and diseased vessels (Figure 2F).

### IFNγ-responsive integrated genes are characterized by STAT1-PU.1 co-binding in combination with increased histone methylation and acetylation and chromatin accessibility

To assess the transcription factor binding motifs, we scanned PU.1 and STAT1 binding sites in specific samples treated with IFNγ (Figure 3A). We detected PU.1 motif and GAS and ISRE motifs as binding sites for PU.1 and STAT1, respectively, in the promoters of treated samples. Besides, it appears that STAT1 binding sites are more enriched in GAS than ISRE.

**Figure 3.**
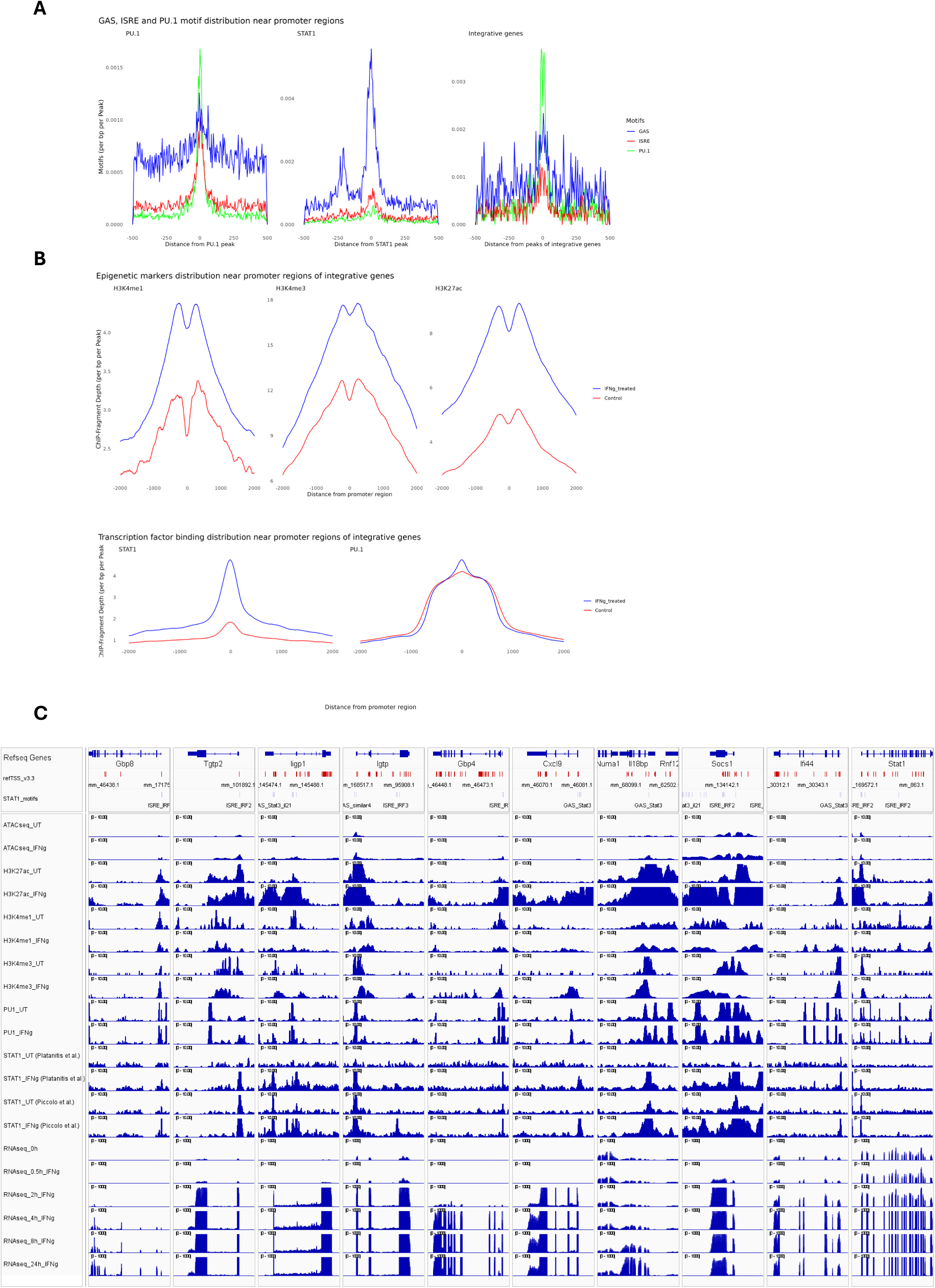
The epigenetic and binding profiles of IFNγ-responsive integrated genes. (A) The enrichment of PU.1 motif and GAS and ISRE motifs as binding sites for PU.1 and STAT1, respectively, in PU.1(left), STAT1(middle) datasets and in the integrative gene peaks (right). (B) The methylation, acetylation and transcription factor binding profiles of integrative genes near the promoter regions. (C) The transcriptional start sites, GAS and ISRE motifs coordinates as well as epigenetic, transcription factor binding and expression profiles of 10 integrative genes were visualized using Integrative Genomics Viewer (IGV). To ensure the consistency of STAT1 binding pattern in response to IFNγ, we showed STAT1 peaks from two separate studies.

To further characterize the integrative genes, we then focused on the promoter regions of these genes. We found the localized distribution of STAT1 and PU.1 motifs surrounding the transcription start site (TSS) of integrative genes. As seen in figure 3A, GAS, ISRE and PU.1 motifs are highly enriched in the promoters of these genes. Moreover, the differential methylation (H3K4me1, H3K4me3) and acetylation (H3K27ac) along with differential binding for STAT1 were markedly prominent in integrative gene promoters in response to IFNγ treatment (Figure 3B). However, as mentioned earlier, PU.1 does not show any differential binding to IFNγ. To obtain a multi-omic perspective of the integrative genes, we prepared an Integrative Genomics Viewer (IGV) snapshot of 10 integrative genes (Figure 3C). As shown in upper tracks, we used refTSS (Abugessaisa et al., 2019), an annotated reference dataset for transcriptional start sites (TSS) in mouse, to identify TSS. Besides, we determined the genomic coordinates of GAS and ISRE motifs which are recognized and bound by STAT1, particularly under IFNγ treatment. It is evident that chromatin is physically accessible for the binding of STAT1 and PU.1, particularly near the promoter regions surrounding TSS, upon exposure to IFNγ. Additionally, there is a distinguished chromatin acetylation which is highly enriched in STAT1 binding sites. On active genes, H3K4me1 and H3K4me3 often co-occur, with H3K4me3 concentrated near the TSS and H3K4me1 flanking these regions (Cheng et al., 2014; Tian et al., 2011), as seen in figure 3C. Besides, we also observed STAT1 gene follows the above-mentioned criteria and shows epigenetic and transcriptional activation. Altogether, it seems that prior to IFNγ treatment the chromatin is in a poised state, characterized mainly by epigenetic modifications including Histone H3K27 acetylation and methylation marks. But upon IFNγ exposure, there is a surge in chromatin openness, increased histone methylation and acetylation along with STAT1-PU.1 co-binding that facilitates the transcriptional activation of STAT1-dependent genes.

### Tracing the expression profile of integrative genes in Atherosclerosis: Macrophage subtype-dependent expression of STAT1-target genes in human atherosclerotic plaques

To better-understand the role of STAT1-dependent integrative genes in a cell specific manner in the context of human atherosclerosis, we analyzed two human datasets with atherosclerotic lesions and one dataset from non-lesion coronary arteries (see Table S1 for the detailed sample description). We integrated these three single cell studies, followed by cell type annotation using a combination of automated and manual approaches, consisting of 40689 cells (Figure 4A; Table S4). The rapid advances in single-cell technologies have facilitated the identification of diverse macrophage subtypes. These subsets express specific markers for pro-inflammatory macrophages (TNF, CXCL2), foamy anti-inflammatory macrophages (TREM2, CD9) and resident-like macrophages (FOLR2, CBR2), found in atherosclerotic plaques in both human and mice (Cochain et al., 2018; Colin, Chinetti-Gbaguidi, & Staels, 2014; Willemsen & de Winther, 2020). In this analysis, we detected foamy macrophages, Monocytes, inflammatory macrophages, tissue resident macrophages and conventional dendritic cells(cDC) in both non-lesion and lesion groups. This observation was also confirmed in other studies (Mosquera et al., 2023; Vallejo et al., 2021). Comparing lesion vs non-lesion groups showed dynamic changes among various macrophage subtypes. For instance, the number of foamy macrophage, monocyte and tissue resident macrophages increased in lesion group (Figure 4B-C). However, an increased proportion of inflammatory macrophages in non-lesion group is due chiefly to the collection of samples from the patients with dilated cardiomyopathies but with no atherosclerotic lesions (Hu et al., 2021).

**Figure 4.**
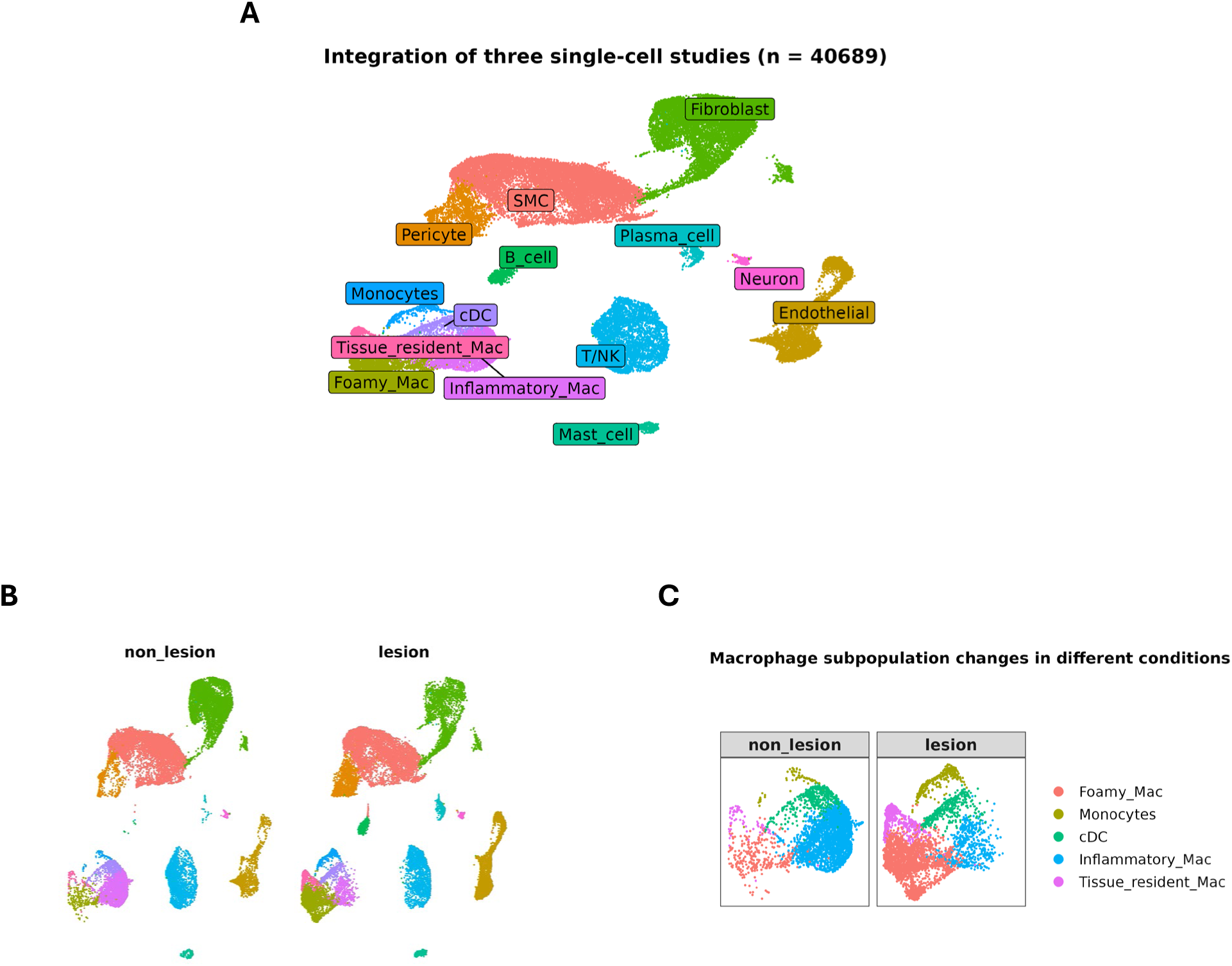
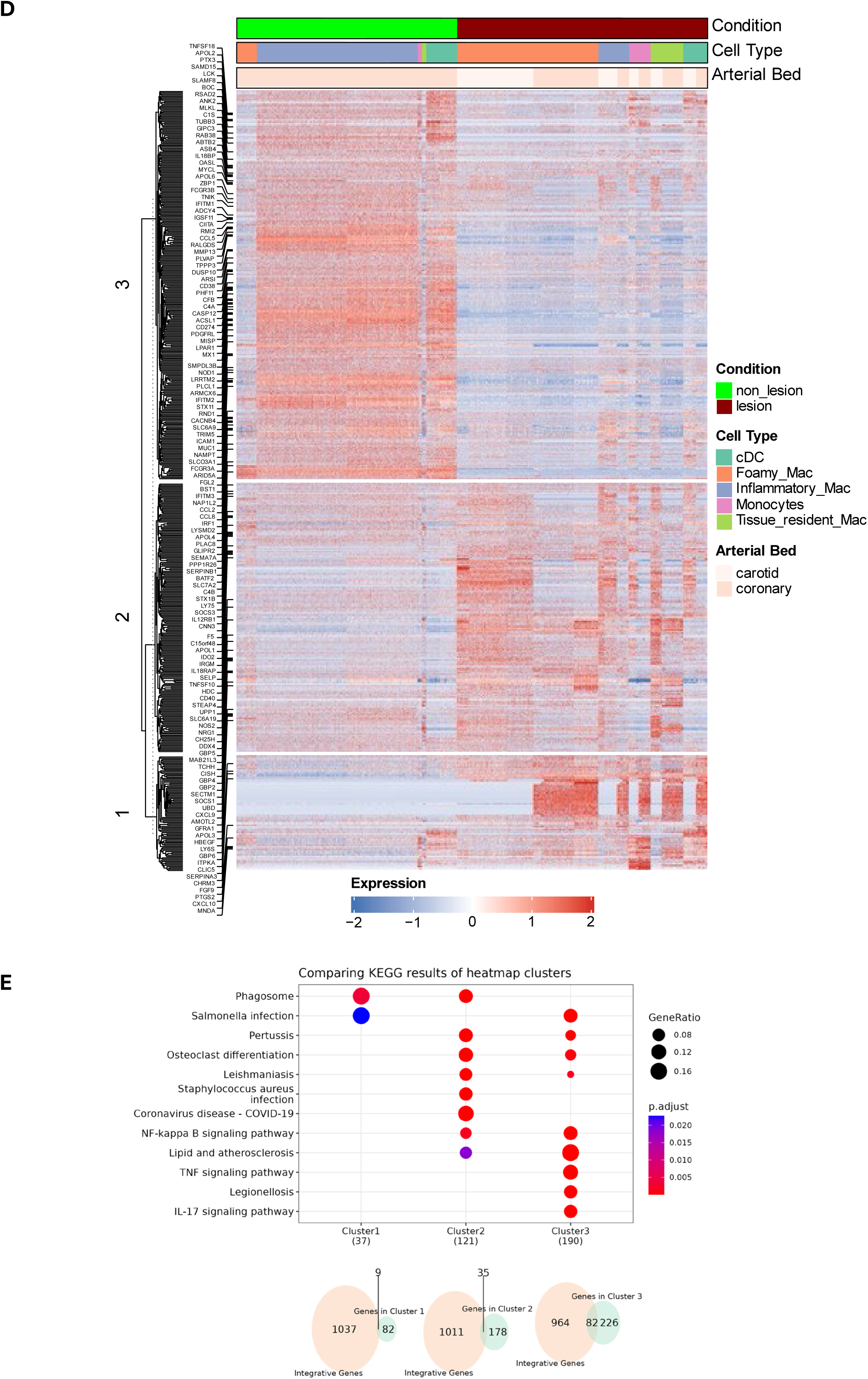
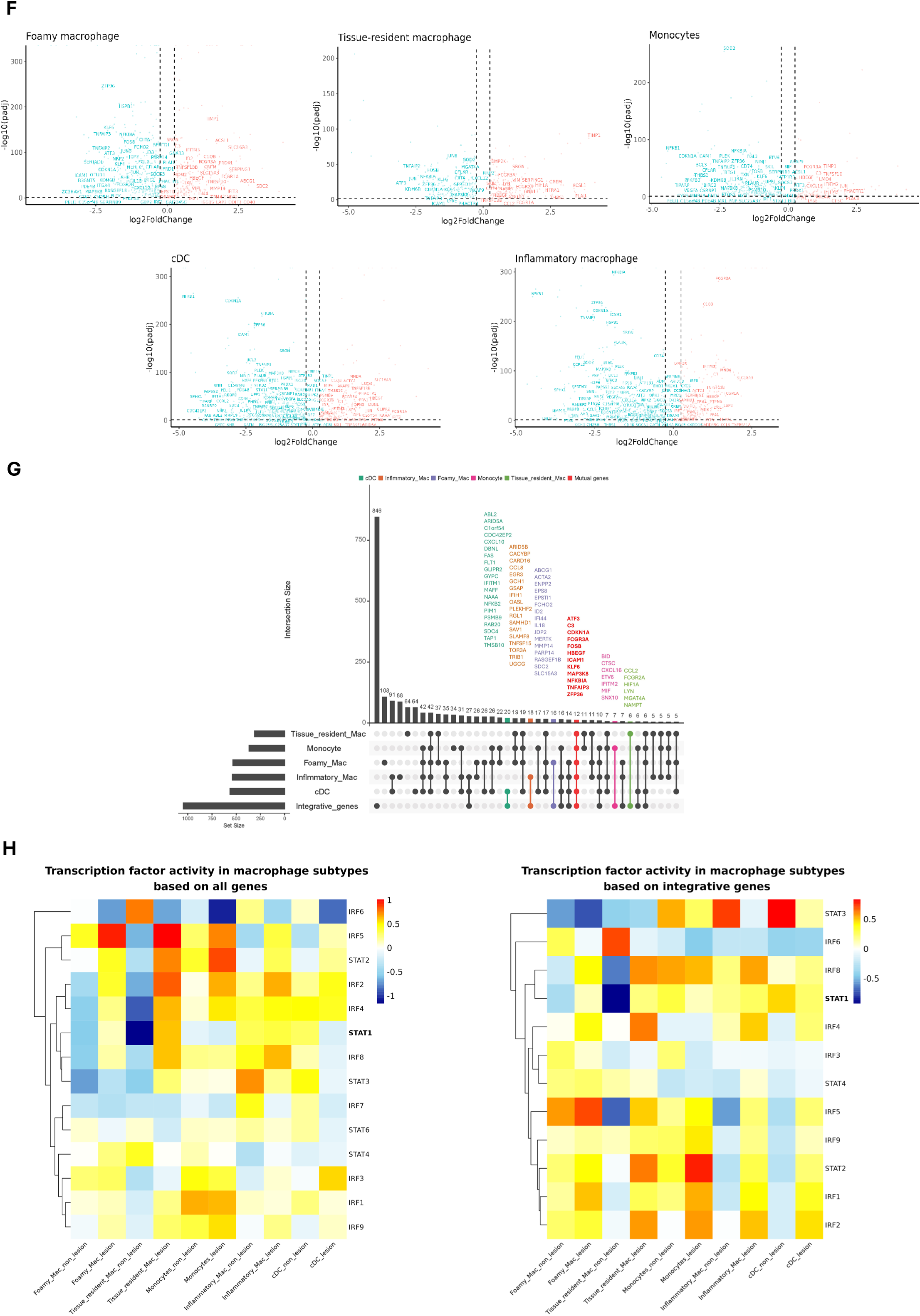
Macrophage subtype-dependent expression of STAT1-target genes in human atherosclerotic plaques. (A) UMAP projection of the three single-cell RNA-seq studies with major cell annotations (See Table S1 for sample details). (B) Comparing lesion vs non-lesion groups showed dynamic changes among various macrophage subtypes. The number of cells in non-lesion and lesion group were 20666 and 20023, respectively. (C) The number of foamy macrophage, monocyte and tissue resident macrophages increased in lesion group. (D) The heatmap representing the expression profile of differentially expressed genes (non-lesion vs lesion) in macrophage population with respect to cell type and arterial bed. The hierarchical cluster analysis generated three distinct clusters. The STAT1-target genes were shown on the left side. (E) KEGG pathway analysis of each cluster showed cluster-specific signaling pathways. (F) The expression pattern of differentially expressed genes (lesion vs non-lesion) in various macrophage subtypes. The STAT1-target genes were labeled in each dot plot. |log2FoldChange| >= 0.25 was used as the cut-off; FDR < 0.05. Red and blue colors represent up-regulated and down-regulated genes, respectively. (G) UpSet plot showing cell type-specific integrative gene sets. The sets on the left (except for integrative genes) indicate the number of cell type-specific differentially expressed genes. The red-colored set shows the unique STAT1-dependent genes present in all macrophage subtypes. (H) The assessment of transcription factor (TF) activity of STAT and IRF family in diverse macrophage subtypes based on all genes (left) or integrative genes (right). For comparison purposes, STAT1 is bolded. The hierarchical clustering on the left side shows TFs with similar activities.

To get a holistic view of the expression pattern of differentially expressed genes (non-lesion vs lesion) in macrophage population, we prepared a heatmap showing the expression pattern with respect to cell type and arterial bed (Figure 4D; Table S5). We then performed hierarchical cluster analysis using euclidean method, generating three distinct clusters. Interestingly, clusters 1 and 2 unveiled arterial bed-specific gene expression pattern with genes in cluster 1 predominantly upregulated in coronary tissue. We next performed KEGG enrichment analysis on each cluster (Figure 4E). The result showed some specific pathway signatures, namely Cluster 1 genes were primarily involved in phagocytosis whereas genes in cluster 2 and 3 were driving inflammatory response and were involved in lipid metabolism. We also assessed the expression profile of differentially expressed genes (non-lesion vs lesion) in vascular smooth muscle cell (VSMCs) population (Figure S1A). Hierarchical clustering generated three clusters with arterial bed-specific gene expressions. For example, cluster 1 genes showed prominently coronary-restricted gene upregulation in lesion group, as also seen in macrophage population. Moreover, it seems that genes in each cluster governing specific signaling cascades, as cluster 1 genes are involved in immune response whereas genes in clusters 2 and 3 are exclusively driving cytoskeleton changes and contraction (Figure S1B). Altogether, it highlights cell-type-specific activities connected to macrophage and VSMCs-selected gene clusters.

We then focused on the expression pattern of STAT1-dependent integrative genes in various macrophage cell types (Figure 4F). Among these cells, inflammatory macrophages, cDC and monocytes showed considerable gene downregulation. Next, we extracted STAT1-dependent integrative genes that were differentially expressed in all of macrophage subtypes (Figure 4G). These genes include ATF3, C3, CDKN1A, FCGR3A, FOSB, HBEGF, ICAM1, KLF6, MAP3K8, NFKBIA, TNFAIP3 and ZFP36. ATF3, as a key transcription factor is involved in regulation of inflammation and lipid metabolism in macrophages (B. Wang et al., 2022). Buono and colleagues reported that disrupted C3 (Complement C3) affects atherosclerosis progression (Buono et al., 2002). Cyclin-dependent kinase inhibitor CDKN1A is involved in inducing cellular senescence in macrophages, contributing to atherosclerosis progression by releasing pro-inflammatory factors (Vellasamy et al., 2022). NFKBIA is shown to be linked with coronary artery disease in the Chinese population (Lai et al., 2015). ICAM1, MAP3K8 and TNFAIP3 are highly associated with the development of atherosclerosis (Garrett et al., 2017; Kitagawa et al., 2002; Sanz-Garcia et al., 2017). FOSB and HBEGF are reported to be upregulated in atherosclerotic plaques (Kim et al., 2019; Liu et al., 2022). KLF6 appears to be a key regulator of macrophage inflammatory responses in the context of atherosclerosis. It mainly promotes pro-inflammatory activation and gene expression while suppressing anti-inflammatory pathways (Kotlyarov & Kotlyarova, 2023) while ZFP36 is engaged in inhibiting pro-inflammatory gene expression (Zhang et al., 2013).

To obtain more insight into the transcription factor (TF) activity of STAT1 regulating the expression of these integrative genes in various macrophage subtypes, we used DoRothEA collection which is a curated dataset of a TF and its transcriptional targets to infer TF activity from gene expression data (Badia-i-Mompel et al., 2022). We inferred TF activity using either all the genes or only the integrative genes (Figure 4H). Close examination of STAT1 revealed prominently increased TF activity in both foamy macrophages and tissue resident macrophages while cDC subtype showing markedly reduced TF activity in lesion group which appeared to be correlated with prominent gene downregulation in cDC. This pattern was also confirmed when the inference of TF activity was performed based solely on the integrative gene expression. We also examined TF activity of other STAT and IRF family members. For instance, STAT2, IRF9, IRF4, IRF5 and IRF8 showed similar activity compared to STAT1.

### Expression profile of STAT1-target genes in the models of mouse atherosclerosis

We also analyzed a mouse single-cell RNA-seq dataset related to atherosclerosis-prone low-density lipoprotein receptor-deficient mice expressing only apolipoprotein B100 with a high-fat diet for three months (Figure 5A; Table S1). The analysis revealed dynamic changes in cell population particularly in macrophages (Figure 5B). We further annotated macrophage population into ISG-expressing immune cells and non-classical monocytes based on the specific makers (IFIT1, IFIT2, IFIT3, IFIT5, ISG15, CCL3, CCL4, CCL3L3, RSAD2, OASL, CXCL10, IFI15, ISG20) and (CD14, CD16, CD11b, CD68, HLA-DR, CD33, CD11c, CD123, CD15, CD3D, CD3E, CD3G, CD3Z, CD66b, FCGR3A, CDKN1C, LST1, FCER1G, MS4A7, RHOC, S100A8, S100A9, CST3, C1QC), respectively (Ianevski, Giri, & Aittokallio, 2022)(Figure 5C). ISG-expressing immune cells generally display inflammatory characteristics (Goel, Kotenko, & Kaplan, 2021). However, non-classical monocytes often show anti-inflammatory properties (Buscher et al., 2017). It should be noted that due to using different experimental protocols, there would be disparities in detection of all macrophage subtypes (Lin et al., 2019). Furthermore, we identified differentially expressed integrative genes in ISG-expressing immune cells and non-classical monocytes in mouse atherosclerosis model (Figure 5D). We detected 16 macrophage-dependent integrative genes in human and LDLR knockout mouse model (Figure 6A). These genes include Ccl5, Ccrl2, Ctsc, Ddit3, Htra1, Id3, Ifitm3, Jun, Ly6e, Marcksl1, Nfkbia, Nupr1, Plaur, Prdx1, Serping1 and Txn1. The CCL2-CCR2 and CCL5-CCR1/CCR5 chemokine axes are critical for monocyte recruitment and early atherogenesis. Blocking these pathways could be a potential therapeutic strategy for atherosclerosis (Márquez, van der Vorst, & Maas, 2021; Zhang et al., 2022). IFITM1 and IFITM3 are two interferon-induced transmembrane proteins that might play significant roles in the pathophysiology of atherosclerosis, particularly through their involvement in inflammation, endothelial function, and vascular health (Friedlová et al., 2022; Kim et al., 2010). The activation of c-Jun is also linked to the inflammatory processes in atherosclerosis (Sozen et al., 2014). Animal model studies have shown that many cathepsin family genes, including CTSC (Cathepsin C), are highly expressed in atherosclerotic plaques (Wang, Jiang, & Cheng, 2022).

**Figure 5.**
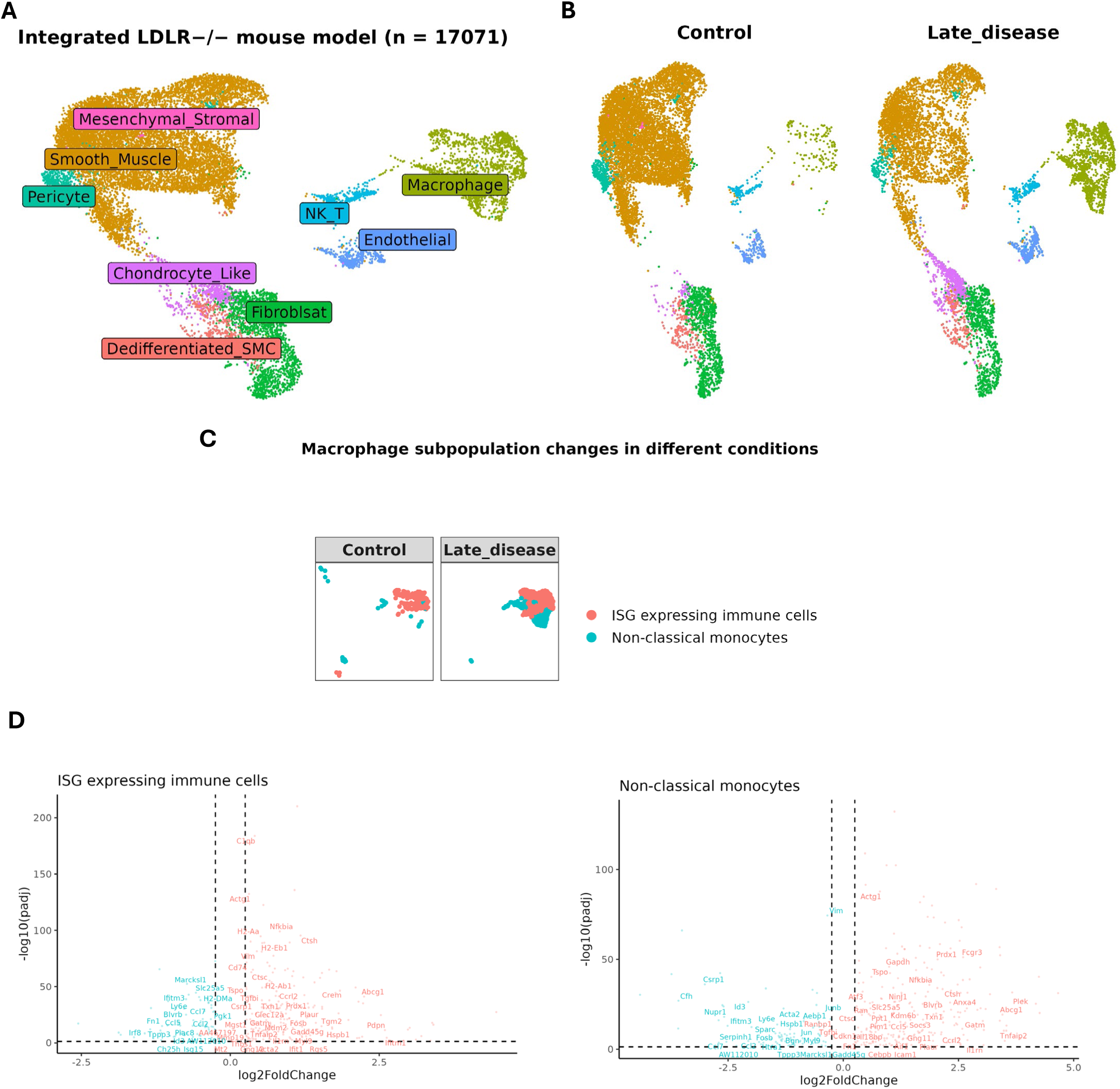
Expression profile of macrophage-dependent STAT1-target genes in LDLR knockout mouse model. (A) UMAP projection of an integrated single-cell RNA-seq study consisting of 17071 cells (See Table S1 for sample details). (B) Comparing late disease vs control groups revealed changes in cell population, (C) particularly in macrophage subtypes namely ISG-expressing immune cells and non-classical monocytes. (D) The expression pattern of differentially expressed genes (late disease vs control) in various macrophage subtypes. The STAT1-target genes were labeled in each dot plot. |log2FoldChange| >= 0.25 was used as the cut-off; FDR < 0.05. Red and blue colors represent up-regulated and down-regulated genes, respectively.

**Figure 6.**
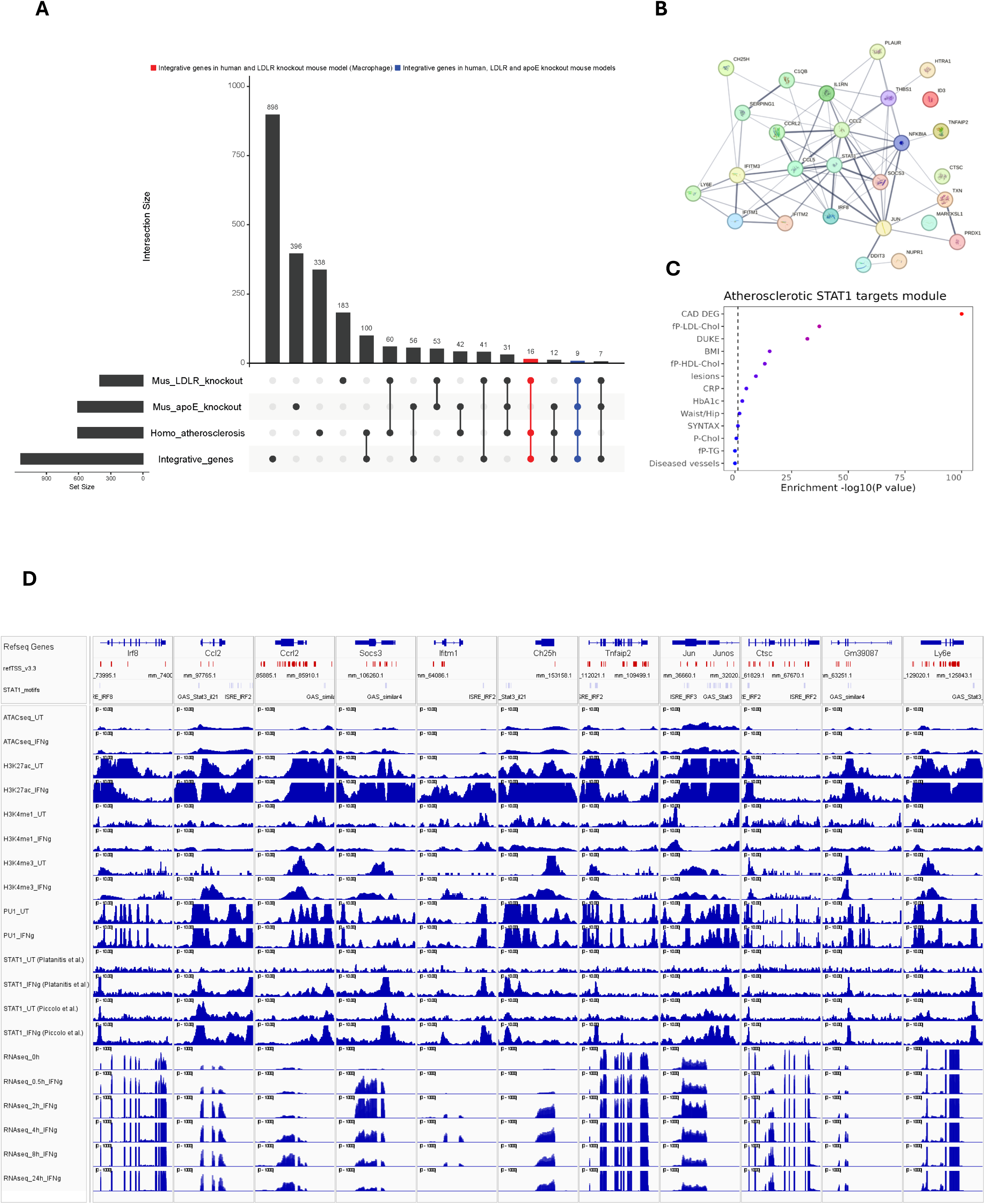
STAT1-dependent gene signature in Atherosclerosis progression. (A) UpSet plot demonstrating the number of integrative genes across mouse and human models. The sets on the left (except for integrative genes) indicate the number of dataset-specific differentially expressed genes. The red-colored set shows the unique STAT1-dependent genes present in all datasets. To obtain 25 signature genes that were differentially expressed in human atherosclerotic plaques and mouse models, the bule and red sets were merged. (B) The protein-protein interaction network of 25 macrophage-dependent STAT1-target gene signature, prepared using STRING database. The thickness of the network edges indicates the strength of data support. (C) The clinical traits that were associated with 25 gene signatures based on STARNET database. The color intensity represents the statistical significance of observed traits. (D) The epigenetic and transcriptomic profiles of 10 signature genes, as described in Figure 3C.

DDIT3 (DNA damage-inducible transcript 3) expression is positively correlated with arterial calcium content and intima-media thickness (IMT) in children with chronic kidney disease (CKD), suggesting it contributes to accelerated arterial calcification and remodeling (Freise et al., 2019). High temperature requirement A1 (HTRA1) is primarily known for its proteolytic activity, which involves the cleavage of various extracellular matrix components. This activity is crucial for maintaining vascular homeostasis and regulating processes such as angiogenesis and vascular remodeling (Dichgans et al., 2023; Ikawati, Kawaichi, & Oka, 2018). Id3 plays a protective role against atherosclerosis. Indeed, Id3-/-ApoE-/- mice develop significantly more atherosclerosis compared to Id3+/+ApoE-/- mice, demonstrating a direct relationship between loss of Id3 and increased atherosclerosis (Doran et al., 2010). Ly6e appears to be a marker of certain macrophage subsets that are enriched in progressing atherosclerotic plaques, suggesting it may play a role in disease progression (Lin et al., 2019). In cardiovascular disease, endothelial polarity proteins like MARCKSL1 help establish endothelial identity and have atheroprotective effects. Endothelial cells secrete extracellular vesicles containing MARCKSL1 in a polarized manner, which can alter monocyte and smooth muscle cell behavior in a compartment-specific way (Raju et al., 2023). NUPR1 is a critical player in the cellular response to stress and oxidative damage. NUPR1’s activation has been linked to increased cardiovascular risk (Fostad et al., 2015; Huang, Santofimia-Castaño, & Iovanna, 2021). Likewise, Txn1, or Thioredoxin-1 helps mitigate oxidative stress within the vascular system (Vogiatzi, Tousoulis, & Stefanadis, 2009). Recent studies have identified PLAUR (Plasminogen activator, urokinase receptor) as an effective diagnostic marker for atherosclerosis lesion progression. Elevated expression levels of PLAUR have been correlated with the severity of atherosclerosis in both human and mouse models (Dai & Lin, 2023). Prdx1 (peroxiredoxin 1) deficiency in macrophages leads to increased susceptibility to oxidative stress and impaired clearance of modified LDL due to defective lipophagic flux, thereby promoting atherosclerosis in apoE-deficient mice (Jeong et al., 2018). Elevated levels of Serping1, also known as C1-inhibitor (C1INH) may indicate a negative prognosis for coronary collateral development, which is important for maintaining blood flow in ischemic conditions (Chen et al., 2022).

We also included a third atherosclerosis data set, in which we performed bulk RNA-seq on aorta from high-fat diet (HFD) fed ApoE knockout mice to identify 763 HFD-upregulated genes (log2FC > 0.5) (Table S6) (Antonczyk et al., submitted).

### Identification of STAT1-dependent gene signature in Atherosclerosis progression

Based on the expression data extracted from the mouse and human models, 9 unique genes including C1qb, Ccl2, Ch25h, Ifitm1, Il1rn, Irf8, Socs3, Thbs1, Tnfaip2 were identified (Figure 6A). The dual role of C1qB in atherosclerosis—both promoting inflammation and providing protective effects—highlights its complexity in disease progression. It has been suggested that the balance between these opposing effects could influence the development and stability of atherosclerotic plaques (Wang et al., 2024; Yang et al., 2021; Zhao et al., 2020). IRF8 appears to play a complex, cell type-specific role in atherosclerosis development, with myeloid IRF8 promoting plaque formation (Louie et al., 2019; Zhang, Gao, et al., 2014). The enzyme cholesterol 25-hydroxylase (CH25H) converts cholesterol into 25-hydroxycholesterol (25-HC), an oxysterol that accumulates in human atherosclerotic lesions, promoting inflammation and plaque instability (Canfrán-Duque et al., 2023). The interleukin-1 receptor antagonist (IL-1Ra), encoded by the IL1RN gene, acts as an important anti-inflammatory brake on IL-1 signaling in the vasculature (Dewberry et al., 2000). SOCS3 (Suppressor of Cytokine Signaling 3) affects macrophage behavior within atherosclerotic plaques. It has been observed that loss of SOCS3 can induce an anti-inflammatory macrophage phenotype, which is beneficial in limiting vascular inflammation and atherosclerosis progression (Taleb et al., 2009; X. Yang et al., 2020). Thrombospondin-1 (TSP-1) is known to modulate inflammatory responses within atherosclerotic plaques. Studies indicate that TSP-1 deficiency leads to increased macrophage infiltration and higher levels of inflammatory cytokines in plaque environments. Specifically, in Thbs1-/- mice, a significant increase in macrophage-induced inflammation was observed, correlating with accelerated plaque necrosis and degradation of elastic lamina due to matrix metalloproteinases (Moura et al., 2008; Stenina & Plow, 2008). Tnfaip2 (Tumor Necrosis Factor Alpha-Inducible Protein 2) enhances inflammatory responses in atherosclerotic lesions. In particular, Tnfaip2 deficiency has been shown to reduce inflammatory cytokine levels and plaque lesions in mouse models of atherosclerosis, indicating its pro-inflammatory role in disease progression (Jin et al., 2022).

We then prepared a combined list of 25 macrophage-dependent STAT1-target gene signature that is differentially expressed in atherosclerotic plaques from human coronary and carotid arteries and aorta from HFD fed LDLR and ApoE knockout mice (Figure 6A). Using STRING database, a protein-protein interaction network was constructed (Szklarczyk et al., 2015). The majority of these 25 genes unveiled functional and physical associations with STAT1 acting as a hub (Figure 6B). Besides, these genes were strongly connected to phenotypic traits such as cardiovascular diseases, cholesterol and lesions (Figure 6C). We also examined if these proteins show chromatin accessibility and are transcriptionally active using bulk datasets related to IFNγ-treated macrophages. As demonstrated in figure 6D, these proteins possess accessible chromatin, exhibit differential acetylation, methylation and STAT1-PU.1 co-binding and are transcriptionally expressed in response to IFNγ.

## Discussion

This study endeavors to provide comprehensive insights into the role of STAT1-mediated IFNγ signaling in atherosclerosis progression through a multi-omics integration approach. By analyzing ATAC-seq, ChIP-seq, and RNA-seq data from IFNγ-treated mouse macrophages, we identified 1139 STAT1-dependent integrative genes that exhibit specific epigenetic and transcriptional characteristics. These genes were further validated in human atherosclerotic lesions and mouse models, revealing significant correlations with LDL cholesterol levels and diseased vessel traits.

The study’s multi-omics approach provides a more detailed understanding of the transcriptional regulation in atherosclerosis compared to previous studies that primarily focused on single-omics data. For instance, while earlier research has established the role of STAT1 in inflammation and atherosclerosis (Lv et al., 2024; Zafar et al., 2021), this study expands on that by demonstrating the integrative epigenetic and transcriptional changes that occur in response to IFNγ.

PU.1 is a transcription factor that plays a crucial role in myeloid lineage specification. The co-binding of STAT1 and PU.1 at target gene promoters aligns with findings from other studies that emphasize the importance of transcription factor interactions in regulating inflammatory gene expression in macrophages (Mancino et al., 2015; Zhao et al., 2022). Our data unveiled that while PU.1 binding was not significantly altered by IFNγ treatment, its co-binding with STAT1 was essential for the transcriptional regulation of the integrative genes. This suggests a collaborative role where PU.1 may help stabilize STAT1 binding or facilitate the transcriptional machinery necessary for gene activation.

STAT1 serves as a pivotal transcription factor in the polarization of macrophages particularly in promoting the M1 phenotype associated with inflammation (Lawrence & Natoli, 2011; Li et al., 2018). In this study, single-cell RNA sequencing of human and mouse atherosclerotic samples revealed dynamic changes in macrophage subtypes, with elevated STAT1 activity observed in both foamy and tissue-resident macrophages. Moreover, the regulation of STAT1 activity is complex and can be influenced by other signaling pathways and transcription factors. We also examined transcription factor activity of other STAT and IRF family members. For instance, STAT2, IRF9, IRF4, IRF5 and IRF8 demonstrated similar activity compared to STAT1. This correlates with a known role of these IRFs in atherosclerosis (Antonczyk et al., 2019; Guo, Callaway, & Ting, 2015; Leipner et al., 2021; Liu et al., 2017; Zhang, Zhu, et al., 2014), suggesting the transcriptional collaboration of STAT1 with IRF family members in these macrophage subtypes.

Our focus on the STAT1-dependent gene signature as potential biomarkers and therapeutic targets aligns with the growing interest in targeting specific signaling pathways related to inflammation in atherosclerosis for therapeutic intervention (Engelen et al., 2022). In this study, we identified 25 macrophage-dependent STAT1-traget genes that were differentially expressed in atherosclerotic plaques from human and mouse models. As explained in the results section, most of these genes were essentially involved in inflammatory response and chemotaxis, lipid metabolism and homeostasis, apoptosis and cellular stress, extracellular matrix and plaque stability and immune regulation. Understanding the roles of these genes provides insight into the complex mechanisms underlying macrophage behavior in atherosclerosis, highlighting potential therapeutic targets for intervention in cardiovascular diseases.

We acknowledge that the study has several limitations that should be properly addressed. Firstly, while the integration of multi-omics data provides a comprehensive view of the transcriptional landscape, it may not fully capture the temporal dynamics of epigenetic modifications over the course of atherosclerosis progression. The use of a specific time point in the analysis may overlook critical changes that occur at other stages of disease development. Secondly, the reliance on murine bone marrow-derived macrophages, while informative, may not completely reflect the complexity of human atherosclerosis, as species differences can influence the response to IFNγ and the resulting macrophage behavior. Additionally, the study primarily focuses on the role of STAT1 in macrophages, potentially underestimating the contributions of other transcription factors, cell types and signaling pathways involved in atherosclerosis. Future studies should aim to address these limitations by incorporating longitudinal analyses and exploring the interactions between various cell types within the atherosclerotic microenvironment.

In conclusion, this comprehensive multi-omics study provides new insights into the transcriptional regulation of atherosclerosis mediated by STAT1-PU.1 co-binding and IFNγ signaling. The identified STAT1-dependent gene signature not only enhances our understanding of disease mechanisms but also highlights potential biomarkers and therapeutic targets for atherosclerosis. Further investigation into the specific roles of these genes in different macrophage subtypes and their interactions with other cell types within the atherosclerotic plaque may lead to the development of more targeted interventions for this complex disease.

## Material and methods

### Macrophage Isolation and differentiation

Bone marrow-derived macrophage samples from C57BL/6 mice of either sex were isolated from femur and tibia by flushing the bones followed by filtration and centrifugation at 500 g. Cells were differentiated for 9–10 days in Dulbecco’s modified Eagle’s medium (DMEM) (Sigma-Aldrich) supplemented with 10% fetal calf serum (FCS), 100 units/ml Penicillin and 100 units/ml Streptomycin (Pen/Strep) (Sigma-Aldrich) and M-CSF (Sigma-Aldrich) on 15-cm Petri dishes. Cells were incubated at 37 °C and 5% CO2 atmosphere. BMDMs were stimulated with IFNγ (100 ng/ml, IF002, MERCK) at specified time points (0, 0.5, 2, 4, 8, 24).

### RNA isolation and RNA-seq library preparation

Total RNA was isolated using TRI-REAGENT (TRI118, MRC) followed by a column-based Total RNA Zol-Out™ D kit (043, A&A Biotechnology) based on manufacturer’s protocol. RNA was quantified using Qubit RNA BR (Broad Range) assay kit (Q10210, TFS) and quality was assessed using Experion™ RNA StdSens Analysis Kit (700-7103, Bio-Rad) according to the manufacturer’s protocol. Only RNA with RNA Integrity Number (RIN) > 9 was considered for library preparation. RNA-seq libraries were prepared in three biological replicates from 1ug of total RNA using NEBNext® Ultra™ II RNA Library Prep Kit for Illumina® (E7770, NEB) together with NEBNext Poly(A) mRNA Magnetic Isolation Module (E7490, NEB) and NEBNext® Multiplex Oligos for Illumina® (E7335, NEB) according to manufacturer’s protocol. Libraries were quantified using Qubit dsDNA HS assay kit (Q32851, TFS) and quality and fragment distribution were examined with Agilent High Sensitivity DNA kit (5067-4626, Agilent Technologies). Sequencing was performed on the NextSeq500 (HighOutput SR75) in Lexogen, BioCenter in Vienna, Austria.

### ApoE KO-based atherosclerosis mouse model

The HFD ApoE KO was essentially performed as described previously (Sanz-Garcia et al., 2017). The experiment was conducted on 16 ten-week-old house mice (Mus musculus) B6.129P2-ApoEtm1Unc/J (purchased from Jacksons Laboratory). Breeding and animal experiments were performed in the animal facility of the Wielkopolskie Centrum Zaawansowanych Technologii (WCZT) in Poznań. All mice work was performed in accordance with the agreement of the Poznan Local Ethical Committee under approval number 16/2019 and 42/2021. Animals were divided into two groups (2x n=8) with mixed sexes. The first group was fed a standard low-fat chow diet (LFD) and the second group of mice was fed a high-fat diet (HFD; High Fat, +7.5 g/kg Cholesterol, Experimental diet, 10.7 % fat, Ssniff S GmbH). After a week of acclimatization and handling, 8-week-old ApoE KO mice were subjected to LFD or HFD for 12 weeks, during which HFD fed mice developed atherosclerotic deposits (Antonczyk et al., submitted).

For RNA isolation, frozen tissues were transferred into Trizol (A&A Biotechnology) and homogenized using a manual Omni tissue homogenizer and dedicated hard tips. All the following steps of RNA isolation were carried out according to Total RNA Zol-Out (A&A Biotechnology) protocol for the rapid purification of ultra-pure total RNA from samples prepared in TRIzol (A&A Biotechnology). RNA-seq library preparation followed the same procedure as for macrophages (see above) (Antonczyk et al., submitted).

### RNA-seq data analysis

The quality of sequencing reads and potential adapter contaminations were evaluated by FastQC (0.12.1) (http://www. bioinformatics.babraham.ac.uk/projects/fastqc/). Low-quality sequences with a Phred score of < 20 were removed by Trim_Galore(0.6.10) (Krueger, 2015). Afterward, the filtered reads were aligned to the mouse genome (GRCm38/mm10) with a fast and efficient spliced aligner tool STAR (2.7.10) (Dobin et al., 2013). FeatureCounts (1.6.2) was employed for the summarization of mapped reads into genomic attributes (Liao, Smyth, & Shi, 2014). Genes with counts lower than 10 at any time points were filtered out. To determine differentially expressed genes (DEG), DESeq2 (1.40.2) package (Love, Huber, & Anders, 2014) in R (4.3.3) was used . The likelihood ratio test (LRT) was implemented to identify genes that respond to IFN treatment over time. False discovery rate (FDR)-adjusted q-values (5% threshold) were calculated by Benjamini–Hochberg procedure. The log2(fold change) FC also was calculated for each gene. Genes with adjusted p-values (padj) less than 0.05 and |log2FC| > 1 were considered as DEGs.

### ATAC-seq data processing

To identify open chromatin regions, the raw sequencing ATAC-seq data was analyzed using nfcore/atacseq pipeline (2.1.2) (Patel et al., 2023). This pipeline is a robust and reproducible method for the processing of ATAC-seq data, which is based on Nextflow (23.04.1). The nfcore/atacseq pipeline includes several stages. Generally, reads were aligned to the mouse genome (GRCm38/mm10) using bwa aligner (0.7.17-r1188) (Li, 2013), followed by peak calling by MACS2 (Gaspar, 2018).

### ChIP-seq data analysis

The raw sequencing ChIP-seq data was analyzed using ENCODE Transcription Factor and Histone ChIP-Seq processing pipeline (https://github.com/ENCODE-DCC/chip-seq-pipeline2) with default parameters as recommended by ENCODE Consortium (Consortium, 2012). Briefly, the sequencing reads were aligned to the mouse genome (GRCm38/mm10) using bowtie2 (2.3.4.3) (Langmead & Salzberg, 2012). Then, duplicates were marked using Picard Tools (2.20.7)(https://github.com/broadinstitute/picard). Peak calling for transcription factors and histones was performed using SPP and MACS2, respectively with FDR threshold set to 0.01. Afterwards, Irreproducible Discovery Rate (IDR) was implemented to identify an optimal number of reproducible peaks between biological replicates, with an IDR score threshold of 0.05.

### Correlation Analysis

The normalized peak files related to ATAC-seq and ChIP-seq data were merged using merge function in bedtools package (Quinlan & Hall, 2010), followed by counting peaks in each datasets using featureCounts (Liao, Smyth, & Shi, 2014) and combining all the count tables into single table for the assessment of correlation using pearson method.

### Identification of differential peaks and integration of datasets

To standardize all sequence alignments from differet datasets (ATAC-seq, STAT1, PU.1, H3K27ac, H3K4me1 and H3K4me3), “Tag Directory” was created using the Homer function makeTagDirectory (Heinz et al., 2010). Then, to find peaks that are differentially enriched between two conditions, the Homer function getDifferentialPeaks was implemented. These normalized differental peaks located in the promoter region region (−3000, 3000) were further selected by ChIPseeker (1.36.0) (Q. Wang et al., 2022), followed by preparing a list of mutual peaks associated with above-mentioned datasets. Next, those peaks were integrated with up-regulated, adjusted p-value (padj < 0.05) genes from our in-house RNA-seq data using BETA tool (1.0.7) (Wang et al., 2013). The upregulated direct target list was selected as integrative genes for downstream analysis.

### Identification of motif instances and distribution of epigenetic markers near the promoter regions

To quantify instances of GAS, ISRE & PU1 motifs in the promoter regions, the Homer function annotatePeaks.pl were implemented on PU.1 and STAT1 peaks in the IFNg-treated group using GAS & ISRE motifs from our previous motif analysis (Sekrecka et al., 2023) and PU.1 motifs from Homer Motif Library (http://homer.ucsd.edu/homer/custom.motifs). To measure the distibution pattern of acetylation and methylation markers and also PU.1 and STAT1 binding sites, Homer function annotatePeaks.pl were employed on the peaks realted to integartive genes.

### Single-cell RNA-seq data analysis

Raw count matrices from each library across the four different studies (mouse and human) were downloaded from GEO and Zenodo (Table S1). The library processing was performed based on the workflow suggested by (Mosquera et al., 2023). Briefly, 17 libraries were processed using Seurat (4.3.0) (Stuart et al., 2019) running in R version 4.3.3. To remove the doublets and ambient RNA, scDblFinder(1.16.0) (Germain et al., 2021) and Celda::DecontX(1.18.1) (S. Yang et al., 2020) R packages were employed, respectively. Then, the decontaminated raw count matrices were furthor filtered to keep the cells that follow 1) >=200 and <=4000 uniquely expressed genes 2) >=200 and <= 20000 UMIs 3) <=10% of reads mapped to the mitochondrial genome 4) <= 5% of reads mapped to hemoglobin genes. Filtered count matrices were normalized using SCTransform (Hafemeister & Satija, 2019). During SCTransform normalization, parameters vst.flavor = “v2” and vars.to.regress = c(“S.Score”,“G2M.Score”) were implemented to take into account for sequencing depth variability and cell cycle variance, respectively. Then, dimensionality reduction of the normalized counts matrix was implemented using Principal Component Analysis (PCA), follwed by applying Uniform Manifold Approximation and Projection (UMAP) non-linear dimensionality reduction using the first 30 PCs.

To intergarte scRNA libraries and remove batch effects, a list of species-specific processed Seurat objects were created, followed by the extraction of 3000 highly variable genes across datasets using SelectIntegrationFeatures. Next, PCA was run across each library using the 3000 variable genes, followed by identification of integration anchors using dimensional reduction method “Reciprocal PCA (rPCA)”, which is an efficient method with respect to the running time and conservation of biological signal. Due to smaller number of mouse datasets, canonical correlation analysis (CCA) were employed instead of rPCA method. The batch-corrected count matrix was then used for PCA dimensionality reduction, creation of the shared-nearest-neighbors (SNN) graph using 30 PCs, and Louvain clustering followed by visualization with UMAP embeddings.

To annotate cell types in a robust manner, we used a blend of automated and manual approaches. For human datasets, we first annotated the integarted data using human cell atalas “Tabula Sapiens (TS)” with a specific focus of Immune and vasculature subset of this atlas (Consortium* et al., 2022). To be consistent with our SCTransformed-integrated datasets, immune and vasculature TS subsets were re-normalized using SCTransform prior to Seurat’s label transfer. We also took advantage of the curated lists of gene markers related to immune and mural cell types in human (Mosquera et al., 2023) and in mouse (Örd et al., 2023) to assess the enrichment score of these genes in our integrated datasets using UCell R pacakge (2.6.2) (Andreatta & Carmona, 2021). Additionally, we also obtained gene markers for each of the SNN-derived clusters using the PrepSCTMarkers and FindAllMarkers functions from Seurat (v4.3.0). Moreover, ScType was implemented for fully-automated cell-type identification based on their comprehensive cell marker database as background information (Ianevski, Giri, & Aittokallio, 2022). Altogether, the cell-annotations of our integrated datasets were finalized using TS atlas, UCell enrichment scores, gene-specific makers for each cluster, ScType predictions and manual confirmation.

### Pseudo-bulk single-cell RNA-seq analysis

For differential expression analyses, we initially identified the 3000 most highly variable genes, followed by retaining these variable genes to accelarte weights computation. We then implemented Zero-Inflated-based Negative Binomial Wanted Variation Extraction (ZINB-WaVE) approach (Van den Berge et al., 2018) using R zinbwave package (1.24.0) to identify excess zero counts and generate gene-and cell-specific weights. The weights were computed taking into account sex, arterial bed and condition as covariates, where applicable. Next, DESeq2 method (Love, Huber, & Anders, 2014) was applied on ZINB-adjusted expression data using single-cell data suitable likelihood ratio test.

### Transcription Factor (TF) activity inference

To infer TF activity, we focused on a specifc list of curated TFs including STAT and IRF familes in DoRothEA R package (1.12.0) (Garcia-Alonso et al., 2019). TFs with high confidience scores were selected and TF activities were then estimated with the R package VIPER (1.36.0) (Alvarez et al., 2016) using the filtered list of regulons and processed Seurat objects which were constructed based on either all the genes or integrative genes. We calculated mean TF activities accros different human macrophage subtypes for either of these seurat objects.

## Data availability

The custom scripts and PU1, GAS and ISRE motifs used in this publication are available on GitHub https://github.com/peculiar97/Atherosclerosis_Multiomics_scRNAseq. The IFNγ-treated macrophage RNA-seq datast and ApoE KO-based atherosclerosis mouse model RNA-seq datset are available through GEO (accession code GSE276418 and GSE270260), respectvely.

**Figure S1.**
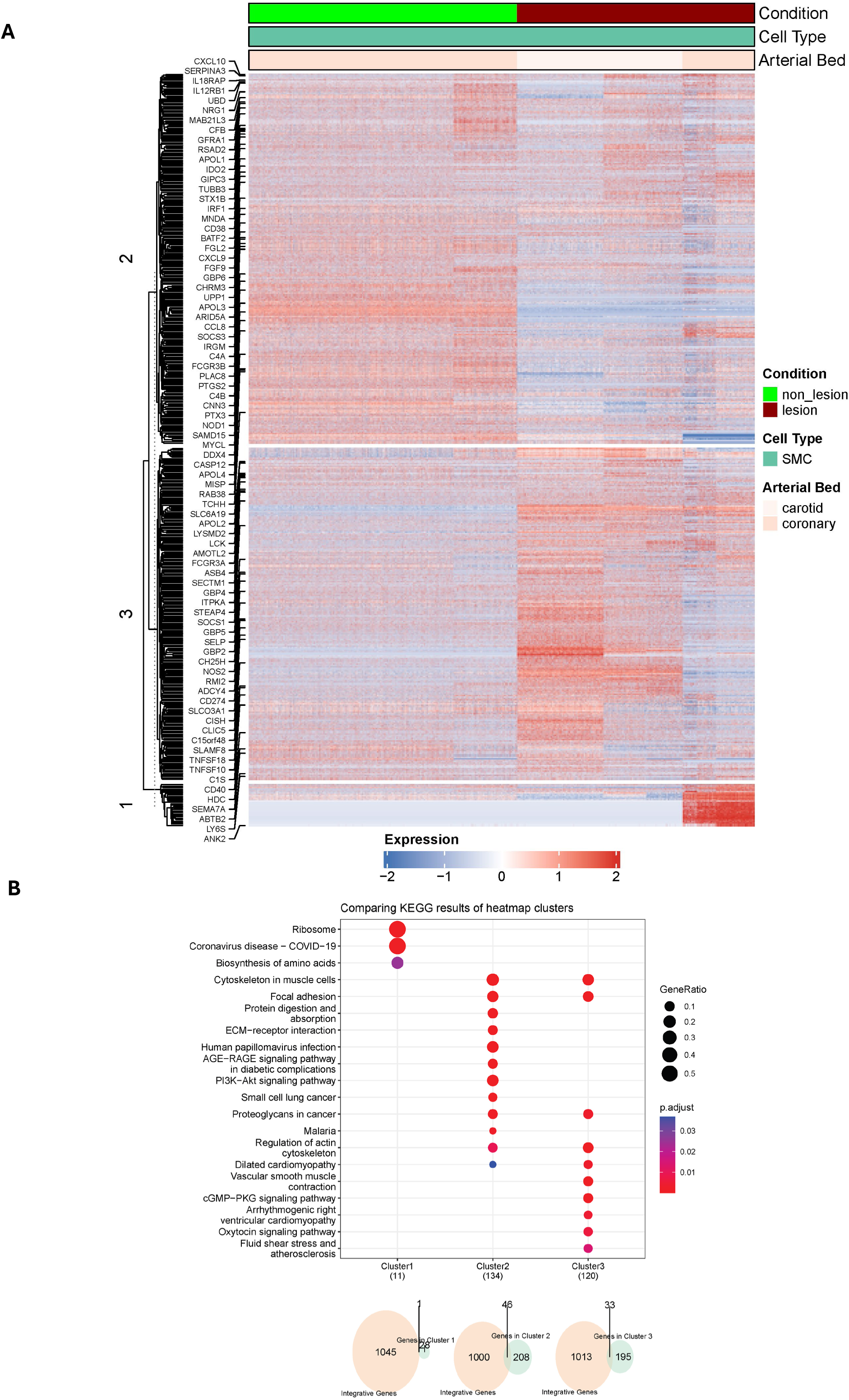
Vascular smooth muscle cell population in human atherosclerotic plaques. (A) The expression profile of differentially expressed genes (lesion vs non-lesion) in vascular smooth muscle cell population. The hierarchical cluster analysis generated three distinct clusters. The STAT1-target genes were shown on the left side. (B) KEGG pathway analysis of each cluster revealed cluster-specific signaling pathways related to muscle activity.

**Table S1.** Public datasets used in this study and sample metadata.

**Table S2.** Differentially expressed genes (DEGs) in IFNγ-treated Macrophages at specified time points (0, 0.5, 2, 4, 8, 24) hours.

**Table S3.** STAT1-dependent integrative gene list.

**Table S4.** Cell type markers were used for the cell annotation of human atherosclerotic plaques.

**Table S5.** The differentially expressed genes in each human macrophage subtype.

**Table S6.** The differentially expressed genes from high-fat diet (HFD) fed ApoE knockout mice.

## Acknowledgements

We thank Dorota Wronka, Anna Karlik, Lukasz Przybyl and Adam Plewinski for their help and advice concerning mouse breeding and tissue collection.

